# Sperm quality metrics exhibit annual fluctuations in a critically endangered amphibian managed under human care

**DOI:** 10.1101/2024.10.07.617017

**Authors:** Allison R. Julien, Isabella J. Burger, Carrie K. Kouba, Diane Barber

## Abstract

The Puerto Rican crested toad (PRCT) is Puerto Rico’s only endemic toad and has undergone rapid population decline within the last 40 years. As a hedge against extinction, captive assurance colonies have been established at several zoological institutions for breeding and reintroduction. However, reproductive output has remained low, despite the use of hormone therapies to attempt to bypass missing abiotic cues and stimulate reproductive behaviors. This low output necessitates a better understanding of natural fluctuations in gamete, specifically sperm, quality in captive individuals. To generate an understanding of natural gametic cycles in captive PRCTs, we administered male PRCTs (n = 86) housed under natural temperature and photoperiod cycles with exogenous hormones monthly for one year. Samples were analyzed for motility and concentration to assess variations in sperm quality by month. Spermiation was successfully stimulated every month, but quality fluctuated; sperm motility was highest in June and July, while sperm concentration was highest in December, January, and March. These results indicate that, while hormones can be utilized to stimulate gamete production in PRCTs year-round, sperm quality is not consistent. Furthermore, the seasonal occurrence of peak sperm production of captive males differed from natural peaks reported for wild PRCTs. Our results illustrate the need for more biologically informed strategies for breeding of at-risk anuran populations.

## 1. Introduction

The Puerto Rican crested toad (*Peltophryne lemur*) is Puerto Rico’s only endemic toad species. Unfortunately, the species has experienced marked declines in the last 40 years, leading to its initial federal listing as “threatened” in 1987 (USFWS 1992) and, following further decline, its global listing as “endangered” in 2020 (IUCN 2021). Cited reasons for this decline are primarily climate change and habitat loss (Burrowes et al. 2004). Puerto Rico has seen increases in both temperature and drought conditions within the last three decades (Larsen et al. 2000; Mote et al. 2017; Gutiérrez-Fonseca et al. 2020). Threats from invasive species are additional stressors; the marine toad (*Rhinella marina*), for example, is a major competitor with *P. lemur* for habitat and prey due its prolific reproductive rates and hardy nature (Joglar et al. 2007; Campbell 2014). Species decline and continued environmental threats have thus led to calls for additional efforts to bolster wild *P. lemur* populations. In response, several zoological institutions have established captive assurance colonies and breeding programs as a means of safeguarding genetic diversity and establishing new wild populations with captive-bred offspring (Clulow et al. 2014).

Currently, the average annual tadpole release from all 18 participating institutions is approximately 42,000 to one reintroduction site; owing to the mortality rates of larval amphibians (Calef 1973; Nolan et al. 2023), these numbers are far below what is necessary to increase and establish new wild populations. Unfortunately, increasing the number of tadpoles for release has been challenging, as many programs have a similar and recurring problem: poor reproductive output. The captive breeding window for the species is large: *P. lemur* reaches sexual maturity as early as 1 year of age in males and 3 years of age in females and has a reported longevity of up to 21 years (Barber 2018). However, due to several factors such as housing and staff limitations, shipping costs, and genetic importance, only two to six pairs of *P. lemur* are bred per institution annually. From these pairings, most institutions report that < 50% of breeding pairs will successfully lay eggs, and in those clutches, fertility rates remain low. Thus, it is crucial that we not only increase the number of successful breeding pairs, but also increase the fertilization rates in each egg clutch.

Challenges to captive breeding are not unique to this species – stimulating reproduction in captive amphibians is difficult due to the reliance of many species on abiotic cues that trigger reproductive behavior and gamete production. These triggers are often species-specific and difficult to replicate (Germano et al. 2009; Kouba et al. 2013; Bronson et al. 2021). When attempts to stimulate breeding behaviors naturally are ineffective, many programs have utilized assisted reproductive technologies (ARTs) to increase fertility rates. The most common of these technologies is exogenous hormone therapy which is used to bypass environmental cues to induce breeding behaviors and gamete production (Kouba et al. 2009; Vu and Trudeau 2016). Breeding programs for the Puerto Rican crested toad have utilized hormones to induce reproductive behaviors since the 1990s with variable success. While protocols have been fine-tuned over the decades, breeding and fertility are not guaranteed and rely on a complex combination of hormone use, brumation, and husbandry practices that mimic seasonal storm conditions (D. Barber, personal communication). Rain and changes in barometric pressure are the most important factors that trigger breeding events for wild *P. lemur* (Cáceres-Charneco et al. 2023). While sporadic breeding has been observed in nearly every month of the year in Puerto Rico, the largest breeding events have occurred in April and May during the rainy season, or August through October during the hurricane season in Puerto Rico (U.S. Fish and Wildlife Service 2016; Keellings and Hernández Ayala, 2019).

Within captive breeding programs, the timing for breeding events has been dictated by suitable environmental conditions for the shipping and rearing of resulting tadpoles and logistical coordination of tadpole transport and release with partners in Puerto Rico for tadpole transport and release. Genearlly captive breeding takes place in June, October, and November (D. Barber, personal communication), which is just outside of the recorded peak breeding months for the species in the wild. It is currently unknown whether or not the decision to breed *P. lemur* outside of historic breeding months has contributed to low reproductive success in these programs with and without the use of exogenous hormones. By monitoring the annual spermatogenic cycle of captive *P. lemur*, we can establish patterns in sperm quality (eg., motility and concentration) and determine how sperm attributes correlate with scheduled breeding events. Thus, the goals of this study were to 1) determine whether spermiation can be successfully induced monthly year-round in captive *P. lemur* using exogenous hormones, and 2) if time of year affects the quality of sperm produced.

## 2. Materials and Methods

### 2.1 Animals

All *Peltophryne lemur* (n = 86) were housed and maintained at the Fort Worth Zoo in Fort Worth, TX for the duration of this study. Males were either captive bred (n = 77) or wild-caught (n = 9). Wild-caught individuals had been housed within the breeding population for 3-5 years at the time of the current study. Average weight and age were 34.5 ± 0.6 g and 4.3 ± 0.2 years, respectively. Weights were established using an electronic benchtop scale. All males were determined to be sexually mature through nuptial pad identification and coloration patterns. Of the 86 animals sampled, 20 had bred during their time in captivity with all 20 having successfully produced offspring. If an animal had been selected for breeding during the study, sperm was not collected for > 2 months before and after breeding to account for any effects of recent breeding history on sperm quality. Animals were housed in mixed-sex groups of three to six individuals in polycarbonate tanks (66 × 36 × 23 cm) filled with water, with ramps allowing for both a dry and aquatic access in each tank. Each tank also contained PVC pipe hides. Diets primarily consisted of crickets gut-loaded with Repashy Superload™ and dusted with an additional vitamin supplement, NektonRep™, and calcium carbonate prior to feeding. All participating animals were determined to be clinically healthy.

Because of the breeding program requirements for the species, *P. lemur* husbandry mimicked seasonal temperature and photoperiod fluctuations. Temperature ranges by season were as follows: 22.7° – 24.4° C in the spring, 24.4° – 30° C in the summer, 22.7° – 24.4° C in the fall, and 22.7° – 28.9° C in winter. A 12-hour day/night photoperiod remained year-round to reflect the 11-13 hour daylight cycles in Puerto Rico. Lighting was provided using UVB fluorescent bulbs above each tank. Husbandry, research protocols, and animal acquisition were approved by the Institutional Animal Care and Use Committee (IACUC) for the Fort Worth Zoo (permit #TE121400-7)

### 2.2 Hormone administration, sperm collection, and analysis

All sperm collections were conducted from October 2021 to July 2023. To stimulate spermiation, male *P. lemur* were treated with a combination of 300 IU human chorionic gonadotropin (hCG; Millipore Sigma, CG5) and 15 µg gonadotropin-releasing hormone (GnRHa; Millipore Sigma, L4513) via intraperitoneal injection delivered in a vehicle of phosphate buffered saline (PBS). These dosages were based on prior research on the species (Burger et al. 2021). Prior to treatment, a baseline urine sample was obtained to confirm that sperm was not being produced naturally. Starting one-hour post-injection, spermic urine samples were collected by holding males above a petri dish until urination occurred, generally within 1 minute. Subsequent collections took place hourly for 8 hours. A total of 10 to 21 males were sampled per month (**Table 1**), and males were re-treated and sampled once every 3-4 months for the duration of the study. Sperm motility and concentration were assessed using an Olympus CX43 phase-contrast microscope. Sperm motility was classified as forward progressively motile (FPM), wherein spermatozoa exhibited forward flagellar movement, motile (M), defined as spermatozoa exhibiting flagellar movement but no forward movement, and non-motile (NM) wherein spermatozoa exhibited no movement. These classifications were reported as a percentage of 100 randomly counted spermatozoa in each sample. Sperm concentration was measured using a hemacytometer (Hausser Scientific #3200). Motility and concentration counts were measured using a manual cell counter.

**Table 1.** Number of male *P. lemur* sampled each month. Weights and age are shown as mean ± SEM.

| Month | n# | Weight (g) | Age (y) |
| --- | --- | --- | --- |
| January | 14 | 33.3 ± 2.1 | 4.0 ± 0.5 |
| February | 17 | 35.7 ± 1.7 | 3.9 ± 0.3 |
| March | 13 | 29.1 ± 1.1 | 4.0 ± 0.3 |
| April | 15 | 38.7 ± 2.1 | 6.4 ± 0.7 |
| May | 15 | 33.2 ± 1.3 | 3.2 ± 0.3 |
| June | 10 | 34.4 ± 2.6 | 5.2 ± 0.4 |
| July | 12 | 32.9 ± 1.9 | 5.3 ± 0.4 |
| August | 11 | 34.9 ± 2.4 | 3.7 ± 1.0 |
| September | 15 | 29.8 ± 1.4 | 1.9 ± 0.4 |
| October | 15 | 33.4 ± 1.8 | 4.7 ± 0.2 |
| November | 21 | 32.7 ± 1.3 | 2.8 ± 0.2 |
| December | 15 | 44.0 ± 0.9 | 6.8 ± 0.3 |

**Table 2.** Results from linear mixed effects models (total motility, forward progressive motility) and generalized linear mixed effects model (concentration).

| Variables | Fixed effects | F statistic (df) | P-value |
| --- | --- | --- | --- |
| **Forward progressive motility (FPM)** | Age | 5.4 (1) | 0.02 |
|  | Weight | 0.36 (1) | 0.55 |
|  | Month | 18.96 (11) | < 0.001 |
| **Total motility (TM)<sup>1</sup>** | Age | 14.36 (1) | < 0.001 |
|  | Weight | 0.34 (1) | 0.56 |
|  | Month | 4.99 (11) | < 0.001 |
| **Concentration** | Age | 0.04 (1) | 0.84 |
|  | Weight | 0.99 (1) | 0.32 |
|  | Month | 8.79 (11) | < 0.001 |

<sup>1</sup>Total motility is calculated as the sum of forward progressive motility (sperm is moving forward) and non-progressive motility (sperm is exhibiting movement but is not moving forward).

### 2.3 Statistical analysis

We used two linear mixed effects models (LMER) to determine the effects of age, weight, and month on 1) sperm forward progressive motility and 2) total motility (TM; FPM + M). Sperm concentration was analyzed using a generalized linear mixed effects model (GLMER) with a Gamma family and link distribution to meet assumptions of normality. Age and weight were included as covariates, and month was included as a categorical fixed effect in every model. Individual toad identity was also added as a random effect. Data are presented as mean ± SEM (standard error of the mean) using the effects package in R, which returns an adjusted mean that incorporates the effects of all variables in the model. We conducted all statistical analyses in R (version 4.3.1).

## 3. Results

### 3.1 Sperm forward progressive motility (FPM) and total motility (TM)

We found that age (p = 0.02), but not weight (p = 0.55), had a significant effect on sperm FPM, with older males producing a lower percentage of sperm with FPM (Fig. 1A). We also found that FPM was affected by the month of collection (p < 0.05). Rates of FPM sperm were higher from December (40.82 ± 4.38%) to July (71.82 ± 4.58%) and lower from July to November (13.85 ± 3.98%) (Fig. 2A).

**Figure 1.**
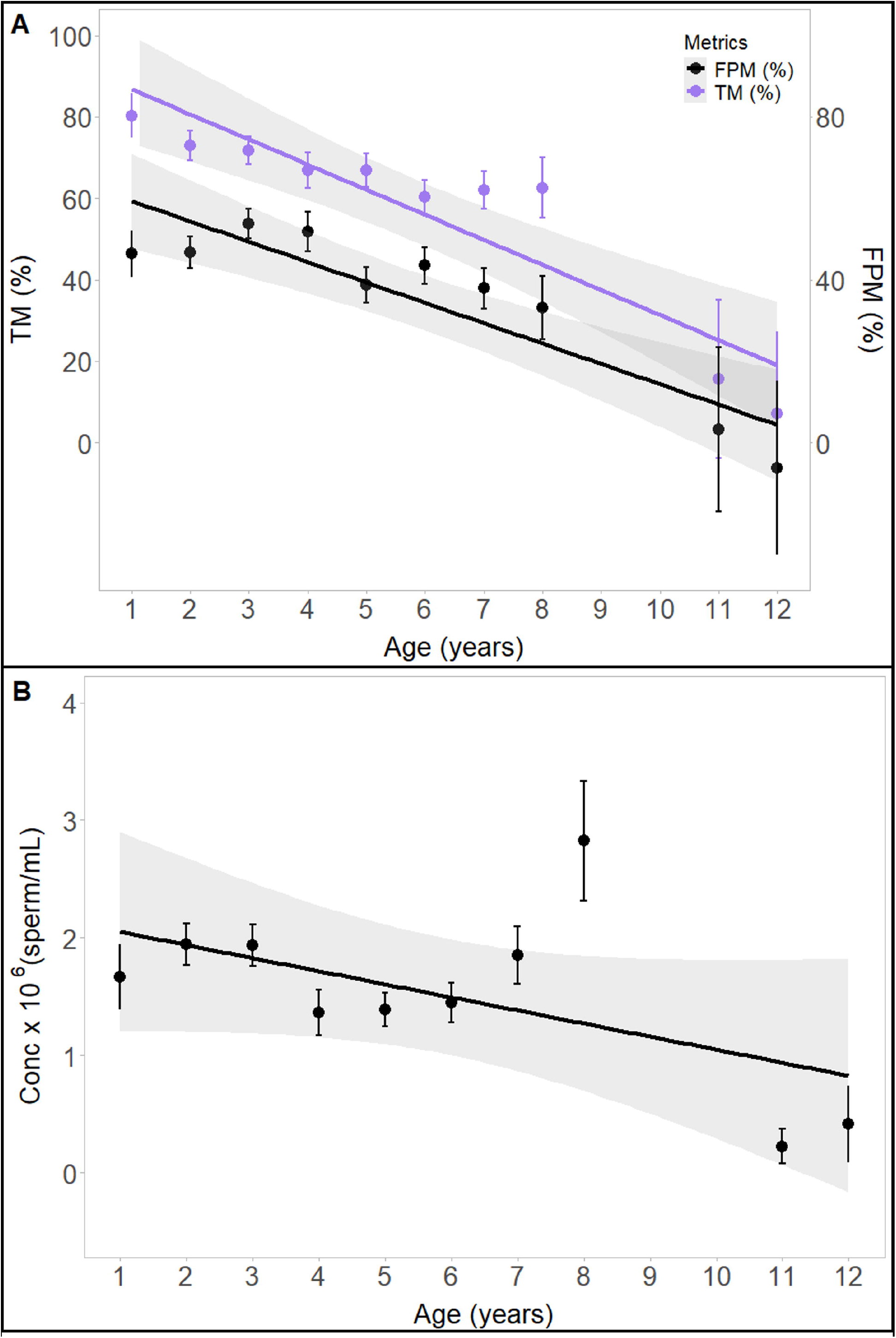
Relationship between age, sperm motility (total motility, TM; forward progressive motility, FPM), and sperm concentration (Conc) for captive *Peltophryne lemur* over a three-year sampling period. Sample sizes for each age at which collection occurred are: 1: n = 15; 2: n = 26; 3: n = 26; 4: n= 14; 5: n = 25; 6: n = 26; 7: n = 15; 8: n = 10; 11: n = 1; 12: n = 1. A) Age had a significant effect on TM and FPM, with both TM and FPM decreasing with increasing age (p < 0.05). B) There was no significant effect of age on sperm concentration (p = 0.84). Figures are partial regression plots that show the relationship of age with TM, FPM, and Conc while accounting for the effects of other variables in the LMER model for TM and FPM and the GLMER for concentration (other variables include weight, month, and individual ID). The trend lines show the trends of FPM, TM, and conc across ages with a 95% confidence interval. Data points and error bars show the mean ± SEM for separate age classes.

**Figure 2.**
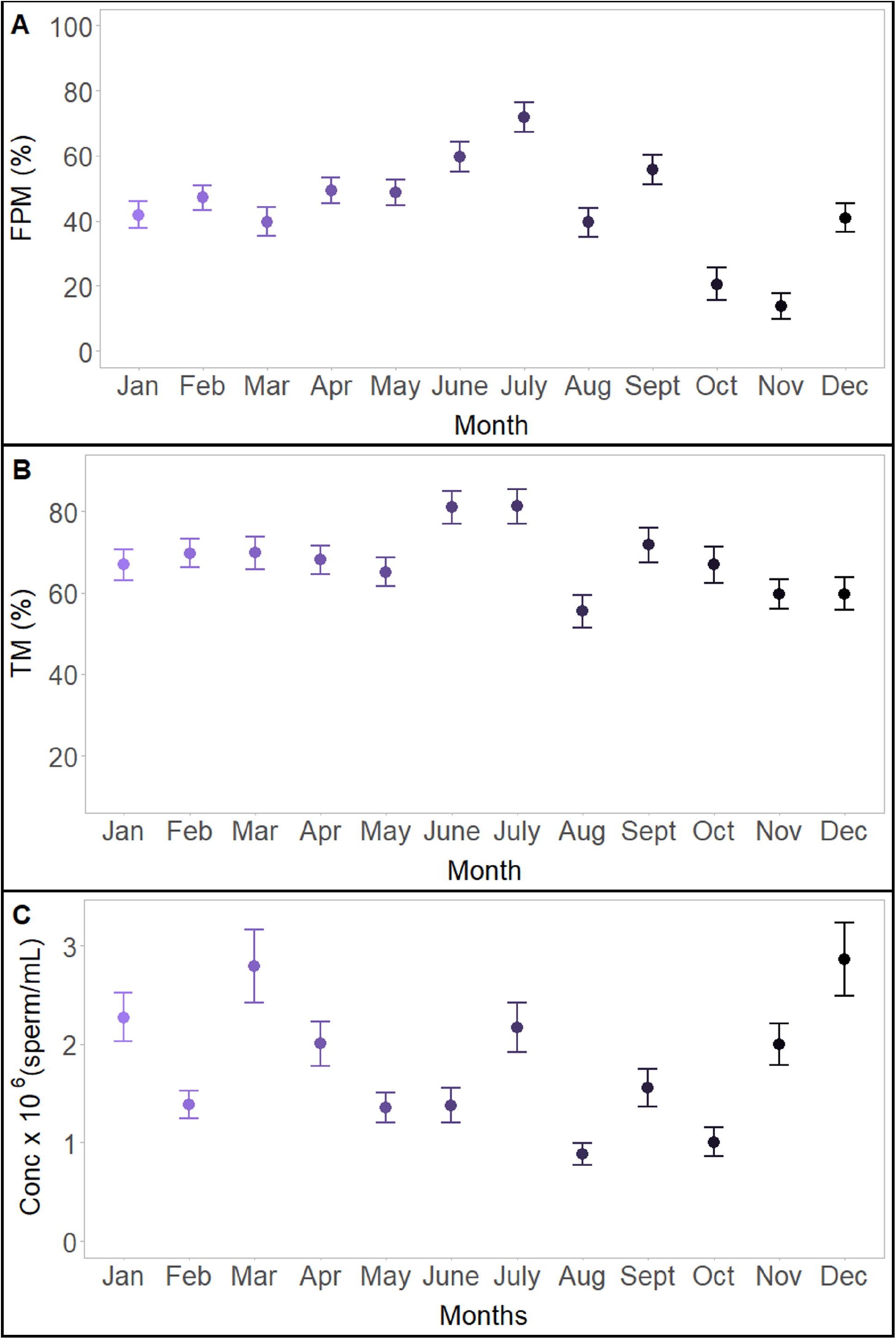
Forward progressive motility (FPM; A), total motility (TM; B) and concentration (Conc; C) of captive *Peltophryne lemur* separated by collection month over a three-year sampling period. Sample sizes for each month at which collection occurred are: Jan: n = 14; Feb: n = 17; Mar: n = 13; Apr: n = 15; May: n = 15; June: n = 10; July: n = 12; Aug: n = 11; Sept: n = 15; Oct: n = 15; Nov: n = 21; Dec: n = 15. Month had a significant effect on all three variables (p < 0.05), with FPM, TM, and Conc varying depending on the month of collection. Figures are partial regression plots that show the relationship of month with TM, FPM, and Conc while accounting for the effects of other variables in the LMER model for TM and FPM and the GLMER for concentration (other variables include weight, age, and individual ID). Data are shown as adjusted mean ± SEM.

The total percentage of motile sperm was also affected by age (p < 0.05) and month (p < 0.05). Similar to FPM, the rate of TM of sperm was lower in older males (Fig. 1A). Regarding variation by month, we found that TM peaked in June (80.95 ± 4.07%) and July (81.27 ± 4.27%). There was no difference in TM from January to May (range 65.10 ± 3.58% – 69.78 ± 3.44%), and TM was lowest in August (55.45 ± 4.03%) (Fig. 2B). Like FPM, there was no effect of weight on TM (p = 0.44).

### 3.2 Sperm concentration

We also found that sperm concentration was significantly affected by the collection month (p < 0.05). Concentration peaked in December (2.86 ± 0.36 × 10^6^ sperm/mL) and March (2.79 ± 0.37 × 10^6^ sperm/mL) and was lowest in August (0.88 ± 0.11 × 10^6^ sperm/mL) and October (1.01 ± 0.15 × 10^6^ sperm/mL) (Fig. 2C). We found no effect of age (p = 0.84) (Fig. 1B) or weight (p = 0.32) on the concentration of sperm released in response to hormone treatment.

## 4. Discussion

One of the benefits of hormone therapies within captive breeding programs is the ability to elicit breeding behavior and gamete production outside of natural breeding seasons or within an imposed time frame (Silla and Byrne 2019). The current study demonstrates that Puerto Rican crested toads can be stimulated to produce sperm year-round with the use of hormone administration. However, our results indicate distinct seasonal trends in sperm characteristics. The highest percentages of motile sperm were observed in June and July, whereas sperm concentrations were highest in December, January, and March. Interestingly, while these two parameters show disparate peaks, occurrence of forward progressive sperm and sperm concentration follow parallel trends (**Fig. 3**), displaying increases in July and September, and decreases in August and October.

**Figure 3.**
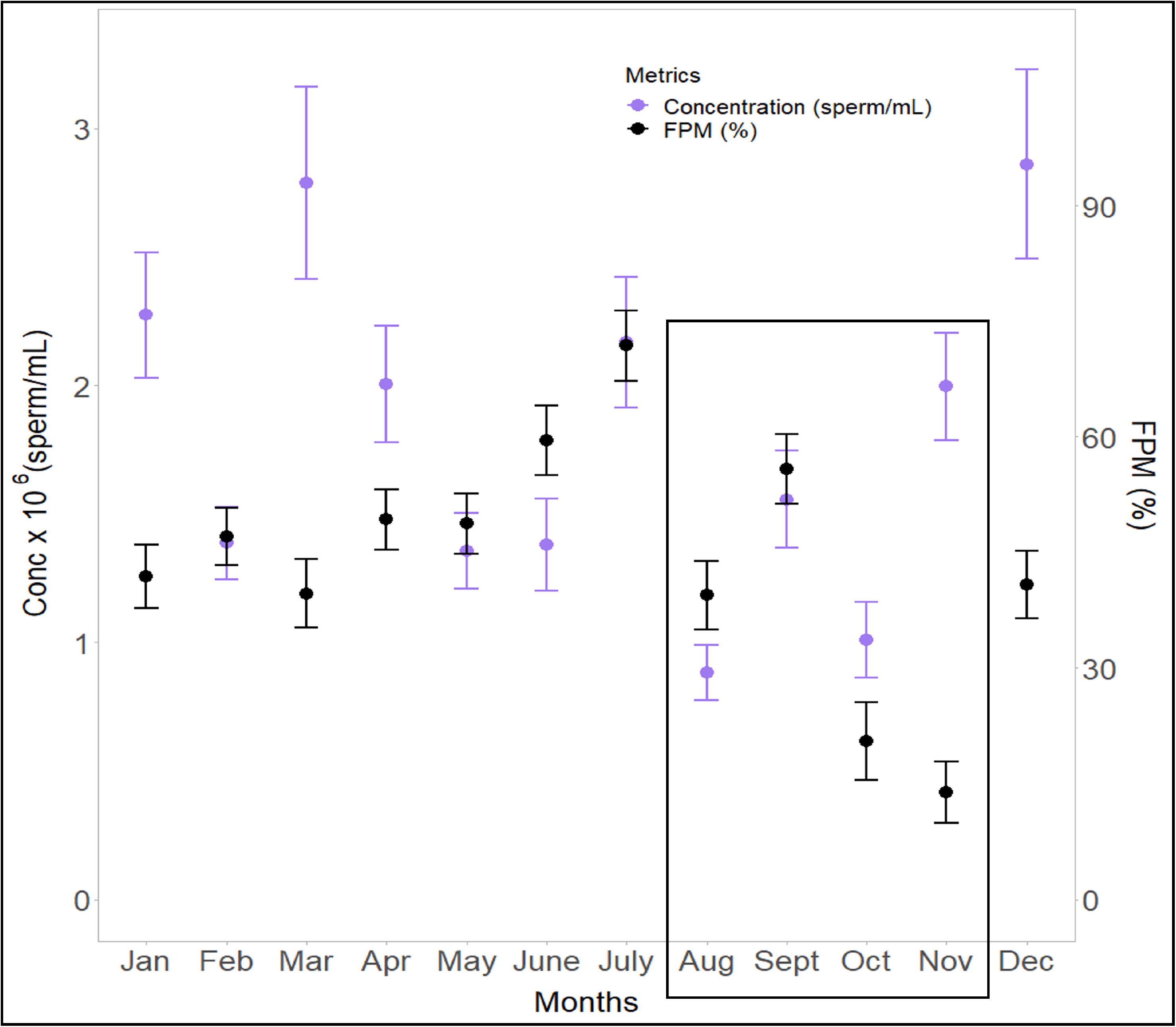
Overlay of forward progressive motility (FPM) and concentration (Conc) fluctuations of captive *Peltophryne lemur* separated by collection month over a three-year sampling period. Sample sizes for each month at which collection occurred are: Jan: n = 14; Feb: n = 17; Mar: n = 13; Apr: n = 15; May: n = 15; June: n = 10; July: n = 12; Aug: n = 11; Sept: n = 15; Oct: n = 15; Nov: n = 21; Dec: n = 15. Months within the natural breeding season for *P. lemur* are indicated by the black box, and arrows indicate months during which captive breeding occurs. The figure is a partial regression plot that show the relationship of month with FPM and Conc while accounting for the effects of other variables in the LMER model for FPM and the GLMER for concentration (other variables include weight, age, and individual ID). Data are shown as adjusted mean ± SEM.

Environmental changes resulting from seasonal differences in precipitation, photoperiod, and temperature fluctuation influence reproductive behaviors in amphibians (Paniagua et al. 1990). For example, many amphibians breed when precipitation levels are high because they depend on specific aquatic conditions for both egg-laying and offspring survival and development (Paniagua et al. 1990). Additionally, spermatogenic cycles in temperate regions often exhibit a discontinuous cycle, where there is high testicular and reproductive activity in the spring and summer, and quiescence in the fall and winter (Paniagua et al. 1990). During quiescent periods, the testes are not responsive to stimulation (Rastogi et al. 2011). Some species of amphibians, however, exhibit a continuous reproductive cycle wherein spermatogenesis and testicular responsiveness occur year-round, although seasonal peaks are still observed (Lofts 1974; Paniagua et al. 1990; Rastogi et al. 2011). While drastic seasonal changes in temperature and photoperiod are common in temperate or colder regions, tropical or equatorial regions, such as Puerto Rico, do not experience such wide variations in these factors annually. However, tropical climates are characterized by changes in precipitation, typically exhibiting dry and wet, or rainy, seasons. Thus, breeding is more strongly correlated with rainy or hurricanic conditions due to shifts in barometric pressure and the formation of freshwater ponds (Matos-Torres 2006). This generally occurs between June and November in Puerto Rico with the greatest peaks in hurricanic activity and associated amphibian breeding, occurring from August-November (Govender et al. 2013; Keellings and Hernández Ayala, 2019).

Unfortunately, changes in barometric pressure and precipitation are extremely difficult to replicate in captivity. In *P. lemur* breeding programs, natural breeding behaviors are rarely observed, even with the use of artificial rain chambers and temperature fluctuations (Barber et al. 2019). At participating institutions, breeding is typically conducted in June and October to coincide with the natural breeding season as well as ensure that environmental conditions are conducive for survival of released offspring. Conversely, in the current study, we found that percentages of motile sperm were lowest in August, October, and November while concentrations were highest in March, November, and December (**Fig. 3**). While the highest percentages of sperm motility and sperm concentration are expected to occur concurrently, we observed contrasting peaks in these two parameters. Possibly, factors affecting sperm quantity (concentration) differ from those affecting sperm quality (motility).

Both sperm motility and concentration are commonly used to assess male fertility, and both factors are heavily influenced by seasonality. Temperature and photoperiod impact spermatogenesis in amphibians through stimulation of the pineal gland, which in turn secretes gonadotropins to stimulate steroid hormone production and subsequent gametogenesis (Paniagua et al. 1990; Mendez-Tepepa et al. 2023). In contrast with other animal taxa, anuran amphibians exhibit a cystic form of sperm release and development, wherein sperm continuously matures in cysts within the testes that burst during the spermiation process, releasing sperm within urine (Mendez-Tepepa et al. 2023). Therefore, the more spermatogenic waves that are initiated through seasonal cues, the higher the concentration of available mature sperm. If males do not breed, mature sperm is not released and instead remains within the testes. Unfortunately, this continuous sperm production and storage can be detrimental to sperm quality over time. For example, in zebrafish (*Danio rerio*), a species observed to mate daily, longer periods of time (eg., > 4 days) without sperm expression resulted in greater concentrations of sperm, but reduced sperm quality, including lower motility rates (Cattelan and Gasparini 2021). The process of spermatogenesis is known to generate high levels of reactive oxygen species (ROS) in the testes through rapid and repeated cell proliferation and renewal (Guerriero et al. 2014; Keogh et al. 2018). High ROS levels can be deleterious to sperm development, survival, and capacity for fertilization (Reinhardt 2007; Keogh et al. 2018; Kowalski et al. 2019). Oxidative stress is associated with the degradation of the cell membrane, increasing membrane permeability and impacting the membrane’s ability to maintain its ionic gradient, which is necessary for flagellar movement (Salisbury and Hart 1970; Reinhardt 2007). In the current study, the disparate peaks of motility (sperm quality) and concentration (sperm quantity) reflect higher rates of cellular degradation due to longer storage times.

The degradation of sperm motility in response to oxidative stress may also be why we found that older males tended to produce lower proportions of motile sperm, but age did not significantly impact sperm concentration (Fig. 3). Disruption of sperm motility is one of the most common consequences of ROS damage arising in ageing sperm (Miller and Blackshaw 1968; Salisbury et al. 2007), and increased ROS production in the testes of older individuals have been well documented (Aitken et al. 1989; Kovac et al. 2013; Chianese and Pierantoni 2021; Anik et al. 2022). The topic of senescence is becoming popular as husbandry methods improve and animals within captive breeding programs have increasing life expectancies or attempts to breed older animals are made when the replenishment of younger animals from the wild is difficult or impossible. While younger animals bred in captivity may be retained within the collection to replenish breeding stock, housing space is limited. Reproductive senescence in amphibians merits further study in the broader context of captive population management.

Since the advent of hormone therapy within captive breeding programs to induce reproduction and gamete expression, natural cyclicity has become a somewhat secondary consideration because of the assumption that it can be overridden. However, while we demonstrate here that sperm can be acquired using hormonal stimulation year-round in *P. lemur*, we also show that there are natural fluctuations in quality. Variation in sperm quantity and quality do not follow presumed historic breeding cycles for the species in the wild, and thus may be impacted by environmental or husbandry conditions while under human care. Asynchronicity between sperm quality and quantity may indicate that human-induced seasonal cycles (i.e., manipulation of light, temperature, moisture, etc.) may not be completely adequate and deleteriously affect the timing of spermatogenesis and sperm storage, perhaps due to the exclusion of understudied factors, such as changes in barometric pressure. In addition, while outside of the scope of the current study, fertilization capabilities should also be examined in reference to seasonality when considering factors affecting captive anuran fecundity. Overall, these results support the incorporation of environmental manipulation of seasonal cues in captive breeding programs for the Puerto Rican crested toad, and other anurans, in order to better coordinate hormone treatment with the species’ natural reproductive cycles. Of course, investigations into cycles for amphibians under human care must be species, and perhaps even population, specific and must be interpreted in the light of wild cycles. Informed collection of fundamental biological data should lead to better reproductive output and cyclicity for programs that face continual limitations of funding and resources.

## 6. Conflict of Interest

The authors declare that the research was conducted in the absence of any commercial or financial relationships that could be construed as a potential conflict of interest.

## 7. Author Contributions

Conceptualization, methodology, writing – original draft preparation, data curation, and investigation, ARJ; Data curation, formal analysis and writing-review and editing, IJB; Conceptualization, writing-review and editing, resources, project administration, and funding acquisition CKK; Conceptualization, writing-review and editing, project administration, and funding acquisition, DB. All authors have read and agreed to the published version of this manuscript.

## 8. Funding

This project was supported by the Fort Worth Zoo and the Institute of Museum and Library Services National Leadership Grants #MG-30-17-0052-17 and MG-251614-OMS-22, and in part by the U.S. Department of Agriculture, Agricultural Research Service, Biophotonics project #6066-31000-015-00D under National Institute of Food and Agriculture Hatch project accession number W4173.

## 9. Acknowledgments

We would like to thank the dedicated zookeeper staff at the Fort Worth Zoo for their care of the animals involved in this study.

## 10. Data Availability Statement

Datasets are available on request: The raw data supporting the conclusions of this article will be made available by the authors, without undue reservation.

## Notes

### Competing Interest Statement

The authors have declared no competing interest.

